# nf-encyclopedia: A cloud-ready pipeline for chromatogram library data-independent acquisition proteomics workflows

**DOI:** 10.1101/2022.09.30.510329

**Authors:** Carolyn Allen, Rico Meinl, Brian C Searle, Seth Just, Lindsay K Pino, William E Fondrie

## Abstract

Data independent acquisition (DIA) mass spectrometry methods provide systematic and comprehensive quantification of the proteome; yet, relatively few open-source tools are available to analyze DIA proteomics experiments. Fewer still are tools that can leverage gas phase fractionated (GPF) chromatogram libraries to enhance the detection and quantification of peptides in these experiments. Here, we present nf-encyclopedia, an open-source NextFlow pipeline that connects three open-source tools—MSConvert, EncyclopeDIA, and MSstats—to analyze DIA proteomics experiments with or without chromatogram libraries. We demonstrate that nf-encyclopedia is reproducible both when run on a cloud platform or a local workstation and provides robust peptide and protein quantification. Additionally, we found that MSstats enhances protein-level quantitative performance over EncyclopeDIA alone. Finally, we benchmarked the ability nf-encyclopedia to scale to large experiments in the cloud by leveraging the parallelization of compute resources. The nf-encyclopedia pipeline is available under a permissive Apache 2.0 license—run it on your desktop, cluster, or in the cloud: https://github.com/TalusBio/nf-encyclopedia.

## 1 Introduction

The ultimate goal of most discovery proteomics methods is to comprehensively detect and quantify the peptides generated from a protein sample. Although data-dependent acquisition (DDA) mass spectrometry methods [1] have long been the workhorse for discovery proteomics experiments, these methods are plagued by missing values between injections onto the mass spectrometer—“runs”—caused by stochastic sampling of the most intense peptide precursor ions during acquisition [2]. In contrast, data-independent acquisition (DIA) mass spectrometry provides run-to-run reproducibility by systematically sampling peptide precursor ions in a cycle of fixed precursor isolation windows [3–5]. However, the reproducibility and theoretical comprehensive nature of DIA methods come at a cost: often DIA isolation windows are large enough to fragment multiple precursor ions simultaneously, resulting in complex tandem mass spectra generated from the fragment ions of multiple peptide precursors. While traditionally the complexity of DIA tandem mass spectra have been challenging to interpret, recent software tools have overcome this barrier [6–11].

One method for increasing proteome identification and coverage with DIA methods has been through the use of chromatogram libraries, built from gas phase fractionated (GPF) pools of samples [10, 12]. In a GPF experiment, a sample is injected multiple times where during each injection narrow precursor isolation windows are cycled over a small precursor mass range [13, 14]. These GPF experiments provide deep, unbiased sampling of the proteome and can be leverage to maximize the information extracted from a quantitative DIA experiment: by collecting a GPF library from a pool of the samples we intend to measure on the same instrument and chromatography column, we gain immense insight into the precise fragmentation of peptides and chromatographic profiles of the peptides we expect to measure. We call these libraries “chromatogram libraries” and they have proven valuable for improving the detection and quantification of peptides in DIA proteomics experiments [10, 15] (Figure S1). Additionally, GPF is easy to implement and not dependent on additional specialty equipment or consumables, unlike offline chromatographic fractionation methods.

DIA experiments with or without chromatogram libraries are most often analyzed by running tools such as EncyclopeDIA, Spectronaut, or DIA-NN on local workstations using their respective graphical user interface. Although this strategy is approachable, it often fails to scale to larger experiments due to the processing time or limited resources of the workstation on which the analysis is performed. Additionally, reproducibility using graphical user interfaces is challenging: documenting where to click is onerous and error-prone, although some software tools provide mechanisms to save and load parameters. When the need to scale beyond a single workstation arises, researchers often resort to using high-performance computing (HPC) clusters and transition to command line interfaces for interacting with their tool of choice. HPC clusters allow for independent processes—individual tasks that must be accomplished in an analysis—to be performed simultaneously by spreading the work across the computers of the cluster—the nodes. This strategy is referred to as “parallelization.” Although HPC clusters can alleviate the problem of scale, they require maintenance, updating, and the constant oversight of a professional. Custom scripts orchestrating series of command line tools on HPC clusters also suffer from reproducibility issues: despite reliably capturing tool parameters, the scripts rarely translate to other computing environments due to their high level of customization.

A growing number of compute-heavy workloads, like DIA proteomics data analyses, are relying on cloud computing resources as an alternative to HPC clusters and single workstations [16]. In the cloud computing paradigm, users essentially rent their own HPC cluster from a cloud provider and are charged only for the resources they use, when they use them. This paradigm allows users to access large networks of computing resources, allowing for parallel processing and the storage of increasingly large datasets [17, 18]. The compute and storage resources available on popular cloud platforms such as Amazon Web Services (AWS), Google Cloud Platform, and Microsoft Azure are virtually unlimited, but typically require expertise to set up and monitor.

Given this diversity of compute platforms, workflow engines such as Nextflow [19] and Snakemake [20] have become indispensable for writing reproducible bioinformatics pipelines. Workflow engines function by separating the pipeline logic—the flow of data through bioinformatics tools—from the compute resources required to execute them. This abstraction allows for portable pipelines to be written in a common language that can be executed on a variety of platforms. Reproducibility across diverse compute platforms is ensured by containerization with tools such as Docker [21] and Singularity [22]. Docker and Singularity images provide a mechanism to encapsulate the required software and all of their dependencies. Analyses that leverage Docker take place within a consistent, “containerized” environment that is identical regardless of the host system. The workflow engine orchestrates the processes needed to complete a bioinformatics analysis, attempting to maximize the efficiency of the available computational resources. Notably, several NextFlow pipelines already exist for analysis of DIA proteomics experiments as part of NF-Core [23,24]; however, these pipelines do not yet support chromatogram library DIA proteomics workflows.

Here, we present nf-encyclopedia, a NextFlow pipeline designed for analyzing DIA proteomics experiments with GPF chromatogram libraries using MSConvert [25], EncyclopeDIA [10], and MSstats [26]. We demonstrate the reproducibility of nf-encyclopedia and assess its quantitative performance. Furthermore, we show that deploying nf-encyclopedia on the cloud can reduce analysis time and enable the efficient analysis of DIA proteomics experiments at scale. Use the nf-encyclopedia pipeline as a power, opensource tool to analyze DIA proteomics experiments with chromatogram libraries at any scale, whether on a local desktop or in the cloud.

## 2 Methods

The nf-encyclopedia pipeline connects three opensource tools—MSConvert [25], EncyclopeDIA [10], and MSstats [26] — in a NextFlow workflow [19] to quantify peptides and proteins from DIA mass spectrometry data files (Figure 1). Furthermore, nf-encyclopedia builds upon configurations and templates provided by NF-Core [23]. Although we designed the nf-encyclopedia pipeline to support the analysis of DIA proteomics experiments with chromatogram libraries, the pipeline also supports the analysis of DIA proteomics experiments with standard spectral libraries.

**Figure 1:**
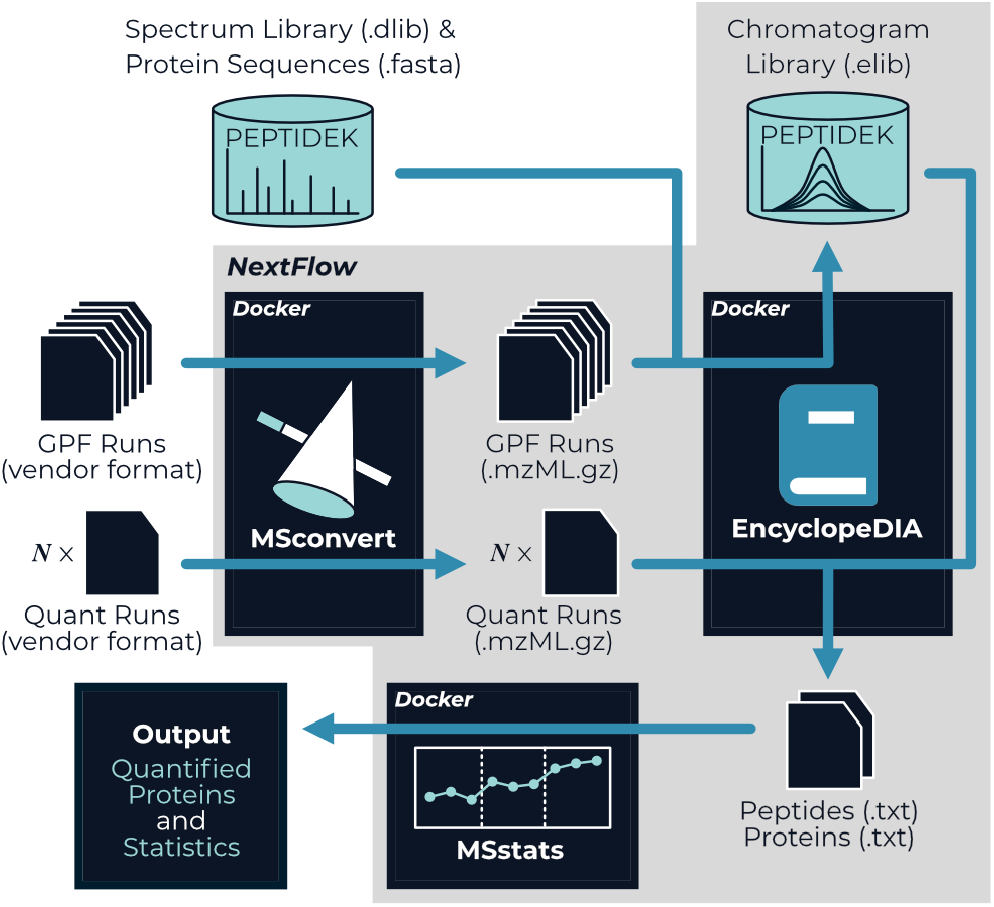
An overview of the nf-encyclopedia pipeline. The mass spectrometry data files are first converted from proprietary vendor formats to a gzipped mzML format using MSConvert. EncyclopeDIA first searches the GPF DIA runs against a provided spectral library to construct a chromatogram library containing retention time and fragmentation information from the same system used to collect the quantitative runs. The chromatogram library is used by EncyclopeDIA to detect and quantify peptides from each quantitative run. Finally, the quantified peptides are processed with MSstats which performs a model-based protein summarization and used to test user-specified hypotheses.

### 2.1 Implementation

The nf-encyclopedia pipeline is written in the Nextflow domain-specific language, which orchestrates the flow of data between the processes that execute MSConvert, EncyclopeDIA, and MSstats. To ensure reproducibility and enable use on common cloud providers, we created a single Docker image containing all of the tools required by the pipeline and it is publicly available on the GitHub Container Registry (https://github.com/TalusBio/nf-encyclopedia/pkgs/container/nf-encyclopedia).Although not required to use nf-encyclopedia, we highly recommend using the provided Docker image for reproducibility and it is enabled by default.

### 2.2 Pipeline Architecture

#### 2.2.1 Input file requirements

The nf-encyclopedia pipeline requires three files: a sample sheet containing the paths to the DIA mass spectrometry data files and whether they comprise a chromatogram library, a FASTA file defining the protein database, and a spectral library in EncyclopeDIA’s DLIB or ELIB format. The sample sheet must contain the path to each file and indicate if each file should be used to build a chromatogram library. It may optionally also define a group that is used to match chromatogram libraries with their associated quantitative DIA runs. This grouping strategy allows for multiple chromatogram libraries to be used in a single analysis, which is critical for longitudinal experiments spanning multiple batches. Additionally, the sample sheet may also specify “conditions” and “bioreplicates” fields to be used by MSstats for model-based quantification, and users can provide a contrast matrix to define the hypotheses to be tested by MSstats. The nf-encyclopedia pipeline is fully compatible with computationally predicted spectral libraries, such as those generated with Prosit [15, 27].

#### 2.2.2 Convert mass spectrometry data files with MSConvert

The mass spectrometry data files provided by the user are first converted to mzML format with gzip and zlib compression using MSConvert, which is able to convert a wide variety of proprietary vendor formats. Additionally, users have the option to let MSConvert demultiplex overlapping DIA isolation windows, a method that improves precursor selectivity with no cost to the duty cycle [12, 28].

#### 2.2.3 Build chromatogram libraries with EncyclopeDIA

If any mass spectrometry data files—typically from gas-phase fractionated DIA runs—are annotated as part of a chromatogram library, nf-encyclopedia will first perform a chromatogram library building step. During this process, each mass spectrometry data file designated as part of a chromatogram library is individually searched by EncyclopeDIA for peptides in the provided spectral library. The search results from individual files are then combined and filtered to achieve a combined 1% false discovery rate (FDR), creating the chromatogram library. Unlike a standard spectral library, the chromatogram library catalogs the retention time, fragment chromatographic peak shapes, fragmentation patterns, and detected interferences for each peptide measured on the system that will be used to quantify peptides and proteins in each sample [10].

#### 2.2.4 Quantify peptides and proteins with EncyclopeDIA

Each mass spectrometry data file is individually searched with EncyclopeDIA for peptides in the chromatogram library, if a chromatogram library was constructed. Otherwise, these files are individually searched for peptides in the provided spectral library. After these searches complete, the pipeline uses EncyclopeDIA to aggregate the individual searches into a single analysis: EncyclopeDIA performs a retention time alignment and “match-between-runs” step to extract quantitative information about each peptide across all runs and filters the aggregate detected peptides and proteins to 1% FDR.

#### 2.2.5 Summarize proteins and calculate statistics with MSstats

Although the peptide and protein quantification tables generated by EncyclopeDIA can be used directly for downstream analyses, nf-encyclopedia also incorporates MSstats to enable model-based protein summarization and to perform statistical hypothesis tests. Users may optionally provide a contrast matrix, which defines the conditions that should be compared in hypothesis tests powered by the linear mixed effects modeling approach used in MSstats.

### 2.3 Assessing Reproducibility

We reanalyzed the pooled HeLa whole-cell lysate samples from the original EncyclopeDIA publication [10] to assess the reproducibility of the nf-encyclopedia pipeline (MSV000082805). This dataset consists of six gas-phase fractionated DIA runs collected using *4-m/z* overlapping isolation windows and three quantitative, single-shot DIA runs collected using *8-m/z* overlapping isolation windows. These runs were demultiplexed using MSConvert to yield 2-*m/z* and 4-*m/z* isolation windows, respectively. We generated a spectral library for the human proteome using Prosit (“Prosit_2020_intensity_hcd” model) for all tryptic peptides with one missed cleavage in 2+ and 3+ charge states at a normalized collision energy of 33 based on a UniProt human proteome FASTA (UP000005640, reviewed, canonical and isoforms, 2022-03 release) [29] with appended contaminant sequences. To assess reproducibility of nf-encyclopedia, we analyzed this dataset in duplicate both on local workstation and in the cloud on AWS Batch and compared the EncyclopeDIA results.

### 2.4 Matrix-matched calibration curves

To test the quantitative performance of the nf-encyclopedia pipeline, we reanalyzed a yeast matrix-matched calibration curve experiment from [30] (PXD014815). This experiment consists of a 14 serial dilutions of yeast peptides (strain BY4741) into a matrix of ^15^N labeled yeast peptides. We compared the limit of detection (LOD) and limit of quantification (LOQ) for peptides and proteins meeting a 20% coefficient of variation (CV) threshold derived from the published EncyclopeDIA results [30] to the EncyclopeDIA and MSstats results generated by our pipeline. Our reanalysis was performed on AWS Batch, using the group functionality of nf-encyclopedia to ensure each of the three replicates were searched against their respective chromatogram libraries. We generated a spectral library for the yeast proteome using Prosit (“Prosit_2020_intensity_hcd” model) for all tryptic peptide with one missed cleavage in 2+ and 3+ charge states at a normalized collision energy of 33 based on the UniProt yeast proteome FASTA (UP000002311, reviewed, canonical, 2021-11 release) with appended contaminant sequences. Lower limits of detection (LOD) and quantification (LOQ) were calculated using the scripts and methods published by [30]. For protein-level figures of merit, our estimates were generated from the protein-level quantitative information summarized by EncyclopeDIA and MSstats.

### 2.5 Benchmarking scalability

We assessed the overall analysis time of the nf-encyclopedia pipeline by reanalyzing increasing numbers of mass spectrometry data files from cerebrospinal fluid of Parkinson’s disease patients (PXD011216) [31]. This dataset contained no gasphase fractionated runs from which to build a chromatogram library, but does contain 289 single-shot DIA runs on a Thermo Q-Exactive HF mass spectrometer using 17 variable isolation windows. We selected random subsets of 3, 5, 10, 25, 50, 100, and 150 raw files and analyzed each subset with the nf-encyclopedia pipeline, in triplicate, on both the cloud using AWS Batch (Amazon Linux 2 host operating system) and on a local workstation (Windows 10 Pro, AMD Ryzen 9 5950X 16-Core processor, 32 GB memory). For these analyses, we used the FASTA file and spectral library that were published with the original study.

### 2.6 Data and Code Availability

All of the mass spectrometry data used in this manuscript is publicly available through ProteomeXchange [32] partner repositories: MSV000082805 on MassIVE, PXD011216 on PRIDE [33], and PXD014815 on Panorama [34]. We used ppx [35] to transfer data from MassIVE and PRIDE to our AWS S3 buckets for analysis. The nf-encyclopedia pipeline is free and open-source software under a permissive Apache 2.0 license, and is available on GitHub where its installation and usage are extensively documented (https://github.com/TalusBio/nf-encyclopedia).

## 3 Results and Discussion

### 3.1 nf-encyclopedia yields reproducible analyses, regardless of platform

A cornerstone of any data analysis pipeline is that it is reproducible: when provided the same data and parameters, the pipeline should yield identical results irrespective of the computational infrastructure or operating system on which it is run. We tested reproducibility of nf-encyclopedia on two platforms, AWS Batch and a local Windows workstation, using the pooled HeLa cell lysate runs from [10]. This dataset consists of six GPF DIA runs from which to build a chromatogram library and three quantitative DIA runs from which to quantify peptides. We analyzed this dataset in duplicate on both platforms using nf-encyclopedia and found that the EncyclopeDIA results were identical in every case: we detected 36,903 peptides and 9323 proteins across the quantitative runs (Figure S2A and B). Furthermore, we found no discrepancies in peptide or protein quantification between replicates or platforms (Figure S2C–E).

Indeed, NextFlow and other workflow managers are designed to facilitate reproducibility and portability between computing platforms by separating the workflow logic from the compute resources. In the case of nf-encyclopedia, we additionally leverage Docker containers to ensure that identical versions of the required software are used on every compute platform and relieve the notorious burden of installing bioinformatics tools. These results confirm that nf-encyclopedia provides a containerized NextFlow workflow that is reproducible between disparate compute environments.

### 3.2 MSstats improves protein-level quantification

Although comprehensive peptide detection and reproducibility are key elements of a DIA proteomics experiment, quantitative performance is where DIA methods excel. To assess the quantitative performance of the nf-encyclopedia pipeline, we reanalyzed a set of matrix-matched calibration curves for the yeast proteome from [30]. These calibration curves consist of serial dilutions of a yeast lysate into a matrix of similar complexity; in this case, a ^15^N-labeled yeast lysate. We use the calibration curves to generate analytical figures of merit—the lower limits of detection (LOD) and quantification (LOQ)—for every quantified peptide. Note that these figures of merit are dependent on both the accuracy and precision of the measurement; hence, a deficiency in either accuracy or precision may lead to a poor LOD and LOQ.

We first compared our reanalysis of the yeast matrix-matched calibration curves from [30] to those reported in the original publication at the peptide level. Our analysis with nf-encyclopedia detected marginally more peptides overall (24,709 in our analysis, as opposed to 24,400 from [12]). Despite differences in this reanalysis—our analysis used EncyclopeDIA v1.34.0 and a predicted spectral library, as opposed to EncyclopeDIA v0.8.0 and Walnut [36] used in the original analysis—we found the distribution of LODs and LOQs for the yeast peptides to be similar (Figures 2A and S3A). This analysis highlights the stability EncyclopeDIA across versions and the ability of our nf-encyclopedia pipeline to reproduce analyses performed outside of the pipeline.

**Figure 2:**
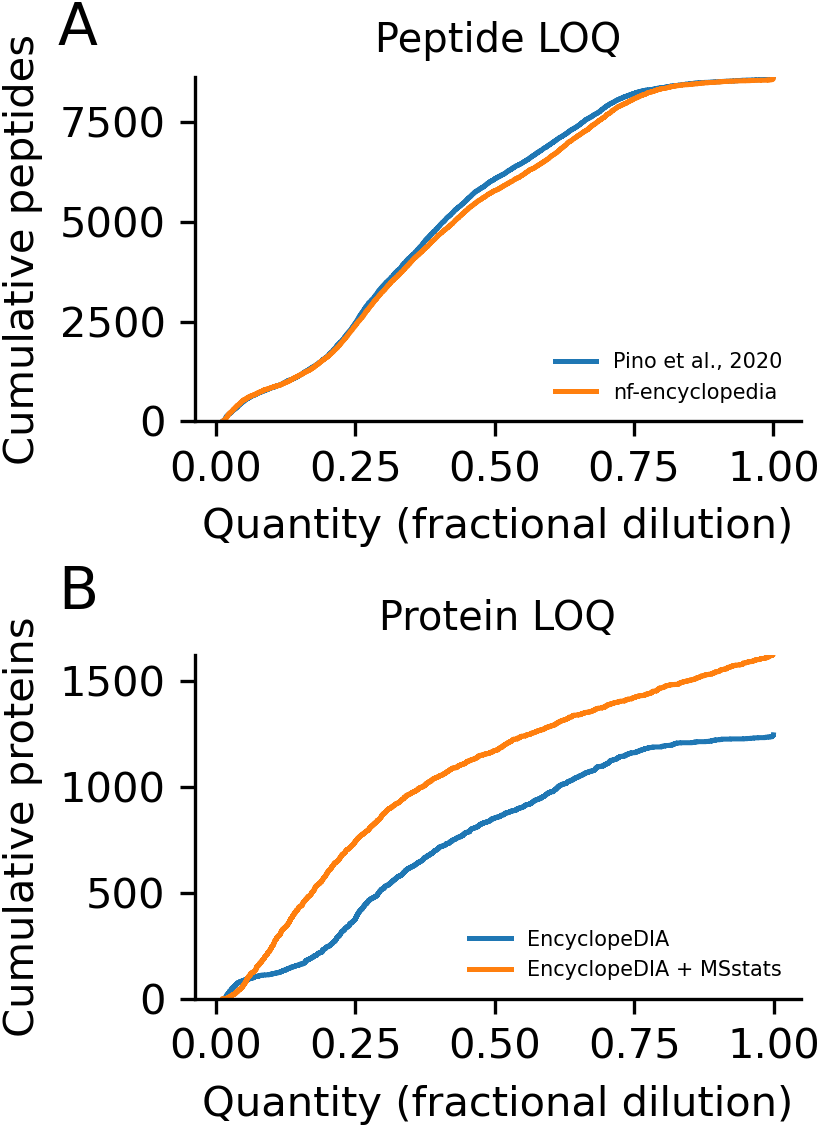
The nf-encyclopedia pipeline maintains and improves quantitative performance. We estimated analytical figures of merit for the peptides and proteins quantified by nf-encyclopedia in a yeast lysate using matrix-matched calibration curves. (A) The LOQ for peptides quantified by nf-encyclopedia followed a similar distribution to the original analysis performed by [30]. (B) MSstats protein summarization improves quantitative performance over EncyclopeDIA alone, shifting the LOQs of proteins toward lower fractional dilutions.

The nf-encyclopedia pipeline provides the option to perform protein-level quantification using MSstats, which we hypothesized may improve quantitative performance over EncyclopeDIA alone. We investigated the LODs and LOQs calculated for every protein using the EncyclopeDIA’s built-in protein summarization method and compared it to those calculated with MSstats protein summarization quantities. Indeed, we found that MSstats generally improved the calculated protein-level LOQs, shifting the LOQ distribution decisively toward lower fractional dilutions (Figure 2B). Conversely, we found that protein summarization with MSstats resulted in slightly poorer LODs in comparison to EncyclopeDIA (Figure S3B); however, this effect is marginal in comparison to the LOQ improvements. As an aside, if we were seeking to minimize the LOD or LOQ of every protein, it would be difficult to improve upon selecting a representative peptide with the lowest LOD or LOQ. However, if we have yet to collect a matrix-matched calibration curve for our samples of interest, these results indicate that the MSstats summarization will yield improved protein-level quantification over EncyclopeDIA alone.

### 3.3 “Infinite” scaling in the cloud

A common issue surrounding the processing of DIA experiments is the ability to scale to large numbers of runs, such as those generated during high-throughput drug screening; hence, we sought to assess the speed at which nf-encyclopedia can process large-scale experiments. One benefit of NextFlow and other workflow managers is that they are readily integrated with cloud providers, such as AWS, Google Cloud Platform, and Microsoft Azure. To benchmark the speed of the nf-encyclopedia pipeline on the cloud, we reanalyzed increasing numbers of DIA run from [31] on our AWS Batch infrastructure and compared wall clock times to the same analyses performed on a local workstation (Figure 3). We found that the workstation was faster for fewer numbers of runs—AWS Batch requires time to recruit suitable compute instances and pull Docker containers before any processing can begin. However, analysis time on the cloud increased at a much lower rate with increasing numbers of runs in comparison to the local workstation: it was able to parallelize independent processes across multiple compute instances, from a pool that is virtually limitless.

**Figure 3:**
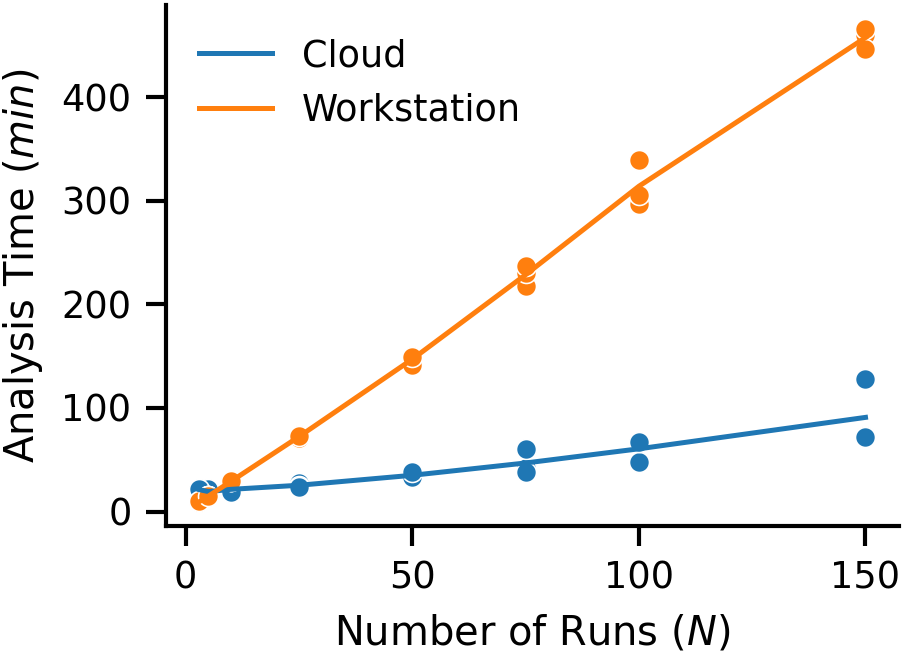
Cloudy with a chance of peptides. The analysis time of the nf-encyclopedia pipeline scales approximately linearly with the number of DIA runs analyzed on a local workstation. However, nf-encyclopedia takes longer to analyze small numbers of runs because the analysis time in the cloud increases at a lower rate as more processes are performed in parallel.

## 4 Conclusions

The nf-encyclopedia pipeline enables the robust processing of DIA proteomics experiments that leverage chromatogram libraries. We demonstrated that nf-encyclopedia is reproducible, maintains high quantitative performance, and scales to large experiments in the cloud. Bioinformatics pipelines written with NextFlow and other workflow engines, such as those maintained by NF-Core [23], provides many of these same advantages for other applications: open-source bioinformatics tools bolster transparency, build community, and are the cornerstone for building reproducible analyses. We anticipate that nf-encyclopedia will become a valuable resource for the proteomics community, connecting open-source tools to provide robust, scalable analyses of “big data” DIA proteomics experiments with chromatogram libraries.

## References

[1] Stahl, D. C., Swiderek, K. M., Davis, M. T., and Lee, T. D. “Data-Controlled Automation of Liquid Chromatography/Tandem Mass Spectrometry Analysis of Peptide Mixtures.” In: Journal of the American Society for Mass Spectrometry 7.6 (June 1996), pp. 532–540.

[2] Webb-Robertson, B.-J. M., Wiberg, H. K., Matzke, M. M., Brown, J. N., et al. “Review, Evaluation, and Discussion of the Challenges of Missing Value Imputation for Mass Spectrometry-Based Label-Free Global Proteomics.” In: Journal of Proteome Research 14.5 (May 2015), pp. 1993–2001.

[3] Venable, J. D., Dong, M.-Q., Wohlschlegel, J., Dillin, A., et al. “Automated Approach for Quantitative Analysis of Complex Peptide Mixtures from Tandem Mass Spectra.” In: Nature Methods 1.1 (Oct. 2004), pp. 39–45.

[4] Gillet, L. C., Navarro, P., Tate, S., Röst, H., et al. “Targeted Data Extraction of the MS/MS Spectra Generated by Data-Independent Acquisition: A New Concept for Consistent and Accurate Proteome Analysis.” In: Molecular & cellular proteomics: MCP 11.6 (June 2012), O111.016717.

[5] Navarro, P., Kuharev, J., Gillet, L. C., Bernhardt, O. M., et al. “A Multicenter Study Benchmarks Software Tools for Label-Free Proteome Quantification.” In: Nature Biotechnology 34.11 (Nov. 2016), pp. 1130–1136.

[6] Tsou, C.-C., Avtonomov, D., Larsen, B., Tucholska, M., et al. “DIA-Umpire: Comprehensive Computational Framework for Data-Independent Acquisition Proteomics.” In: Nature Methods 12.3 (Mar. 2015), 258–264, 7 p following 264.

[7] Demichev, V., Messner, C. B., Vernardis, S. I., Lilley, K. S., et al. “DIA-NN: Neural Networks and Interference Correction Enable Deep Proteome Coverage in High Throughput.” In: Nature Methods 17.1 (Jan. 2020), pp. 41–44.

[8] Röst, H. L., Rosenberger, G., Navarro, P., Gillet, L., et al. “OpenSWATH Enables Automated, Targeted Analysis of Data-Independent Acquisition MS Data.” In: Nature Biotechnology 32.3 (Mar. 2014), pp. 219–223.

[9] Bruderer, R., Bernhardt, O. M., Gandhi, T., Miladinović, S. M., et al. “Extending the Limits of Quantitative Proteome Profiling with Data-Independent Acquisition and Application to Acetaminophen-Treated Three-Dimensional Liver Microtissues.” In: Molecular & cellular proteomics: MCP 14.5 (May 2015), pp. 1400–1410.

[10] Searle, B. C., Pino, L. K., Egertson, J. D., Ting, Y. S., et al. “Chromatogram Libraries Improve Peptide Detection and Quantification by Data Independent Acquisition Mass Spectrometry.” In: Nature Communications 9.1 (Dec. 2018).

[11] Lu, Y. Y., Bilmes, J., Rodriguez-Mias, R. A., Villén, J., et al. “DIAmeter: Matching Peptides to Data-Independent Acquisition Mass Spectrometry Data.” In: Bioinformatics (Oxford, England) 37.Suppl_1 (July 2021), pp. i434–i442.

[12] Pino, L. K., Just, S. C., MacCoss, M. J., and Searle, B. C. “Acquiring and Analyzing Data Independent Acquisition Proteomics Experiments without Spectrum Libraries.” In: Molecular & cellular proteomics: MCP 19.7 (July 2020), pp. 1088–1103.

[13] Yi, E. C., Marelli, M., Lee, H., Purvine, S. O., et al. “Approaching Complete Peroxisome Characterization by Gas-Phase Fractionation.” In: Electrophoresis 23.18 (Sept. 2002), pp. 3205–3216.

[14] Panchaud, A., Scherl, A., Shaffer, S. A., von Haller, P. D., et al. “Precursor Acquisition Independent from Ion Count: How to Dive Deeper into the Proteomics Ocean.” In: Analytical Chemistry 81.15 (Aug. 2009), pp. 6481–6488.

[15] Searle, B. C., Swearingen, K. E., Barnes, C. A., Schmidt, T., et al. “Generating High Quality Libraries for DIA MS with Empirically Corrected Peptide Predictions.” In: Nature Communications 11.1 (Mar. 2020), p. 1548.

[16] Neely, B. A. “Cloudy with a Chance of Peptides: Accessibility, Scalability, and Reproducibility with Cloud-Hosted Environments.” In: Journal of Proteome Research 20.4 (Apr. 2021), pp. 2076–2082.

[17] Halligan, B. D., Geiger, J. F., Vallejos, A. K., Greene, A. S., et al. “Low Cost, Scalable Proteomics Data Analysis Using Amazon’s Cloud Computing Services and Open Source Search Algorithms.” In: Journal of Proteome Research 8.6 (June 2009), pp. 3148–3153.

[18] Langmead, B. and Nellore, A. “Cloud Computing for Genomic Data Analysis and Collaboration.” In: Nature Reviews. Genetics 19.4 (Apr. 2018), pp. 208–219.

[19] Di Tommaso, P., Chatzou, M., Floden, E. W., Barja, P. P., et al. “Nextflow Enables Reproducible Computational Workflows.” In: Nature Biotechnology 35.4 (Apr. 2017), pp. 316–319.

[20] Köster, J. and Rahmann, S. “Snakemake—a Scalable Bioinformatics Workflow Engine.” In: Bioinformatics (Oxford, England) 28.19 (Oct. 2012), pp. 2520–2522.

[21] Merkel, D. “Docker: Lightweight Linux Containers for Consistent Development and Deployment.” In: Linux Journal 2014.239 (Mar. 2014), 2:2.

[22] Kurtzer, G. M., Sochat, V., and Bauer, M. W. “Singularity: Scientific Containers for Mobility of Compute.” In: PLOS ONE 12.5 (May 2017), e0177459.

[23] Ewels, P. A., Peltzer, A., Fillinger, S., Patel, H., et al. “The Nf-Core Framework for Community-Curated Bioinformatics Pipelines.” In: Nature Biotechnology 38.3 (Mar. 2020), pp. 276–278.

[24] Bichmann, L., Gupta, S., Rosenberger, G., Kuchenbecker, L., et al. DIAproteomics: A Multi-Functional Data Analysis Pipeline for Data-Independent-Acquisition Proteomics and Peptidomics. Preprint. Bioinformatics, Dec. 2020.

[25] Chambers, M. C., Maclean, B., Burke, R., Amodei, D., et al. “A Cross-Platform Toolkit for Mass Spectrometry and Proteomics.” In: Nature Biotechnology 30.10 (Oct. 2012), pp. 918–920.

[26] Choi, M., Chang, C.-Y., Clough, T., Broudy, D., et al. “MSstats: An R Package for Statistical Analysis of Quantitative Mass Spectrometry-Based Proteomic Experiments.” In: Bioinformatics (Oxford, England) 30.17 (Sept. 2014), pp. 2524–2526.

[27] Gessulat, S., Schmidt, T., Zolg, D. P., Samaras, P., et al. “Prosit: Proteome-Wide Prediction of Peptide Tandem Mass Spectra by Deep Learning.” In: Nature Methods 16.6 (June 2019), pp. 509–518.

[28] Amodei, D., Egertson, J., MacLean, B. X., Johnson, R., et al. “Improving Precursor Selectivity in Data-Independent Acquisition Using Overlapping Windows.” In: Journal of the American Society for Mass Spectrometry 30.4 (Apr. 2019), pp. 669–684.

[29] The UniProt Consortium. “UniProt: The Universal Protein Knowledgebase in 2021.” In: Nucleic Acids Research 49.D1 (Jan. 2021), pp. D480–D489.

[30] Pino, L. K., Searle, B. C., Yang, H.-Y., Hoofnagle, A. N., et al. “Matrix-Matched Calibration Curves for Assessing Analytical Figures of Merit in Quantitative Proteomics.” In: Journal of Proteome Research 19.3 (Mar. 2020), pp. 1147–1153.

[31] Rotunno, M. S., Lane, M., Zhang, W., Wolf, P., et al. “Cerebrospinal Fluid Proteomics Implicates the Granin Family in Parkinson’s Disease.” In: Scientific Reports 10.1 (Dec. 2020), p. 2479.

[32] Deutsch, E. W., Bandeira, N., Sharma, V., Perez-Riverol, Y., et al. “The ProteomeXchange Consortium in 2020: Enabling ‘Big Data’ Approaches in Proteomics.” In: Nucleic Acids Research 48.D1 (Jan. 2020), pp. D1145–D1152.

[33] Perez-Riverol, Y., Bai, J., Bandla, C., García-Seisdedos, D., et al. “The PRIDE Database Resources in 2022: A Hub for Mass Spectrometry-Based Proteomics Evidences.” In: Nucleic Acids Research 50.D1 (Jan. 2022), pp. D543–D552.

[34] Sharma, V., Eckels, J., Taylor, G. K., Shulman, N. J., et al. “Panorama: A Targeted Proteomics Knowledge Base.” In: Journal of Proteome Research 13.9 (Sept. 2014), pp. 4205–4210.

[35] Fondrie, W. E., Bittremieux, W., and Noble, W. S. “Ppx: Programmatic Access to Proteomics Data Repositories.” In: Journal of Proteome Research 20.9 (Sept. 2021), pp. 4621–4624.

[36] Ting, Y. S., Egertson, J. D., Bollinger, J. G., Searle, B. C., et al. “PECAN: Library-Free Peptide Detection for Data-Independent Acquisition Tandem Mass Spectrometry Data.” In: Nature Methods 14.9 (Sept. 2017), pp. 903–908.

