## Supplemental Figures for "nf-encyclopedia: A cloud-ready pipeline for chromatogram library data-independent acquisition proteomics workflows"

### List of Figures

S1 The GPF chromtagram library workflow. . . . . . . . . . . . . . . . . . . . . . . . . . . . . . . . . . . S-2
S2 nf-encyclopedia is reproducible in the cloud and a local workstations. . . . . . . . . . . . . . . . . . S-3
S3 MSstats yields slightly poorer protein-level LODs than EncyclopeDIA alone. . . . . . . . . . . . . . . S-4

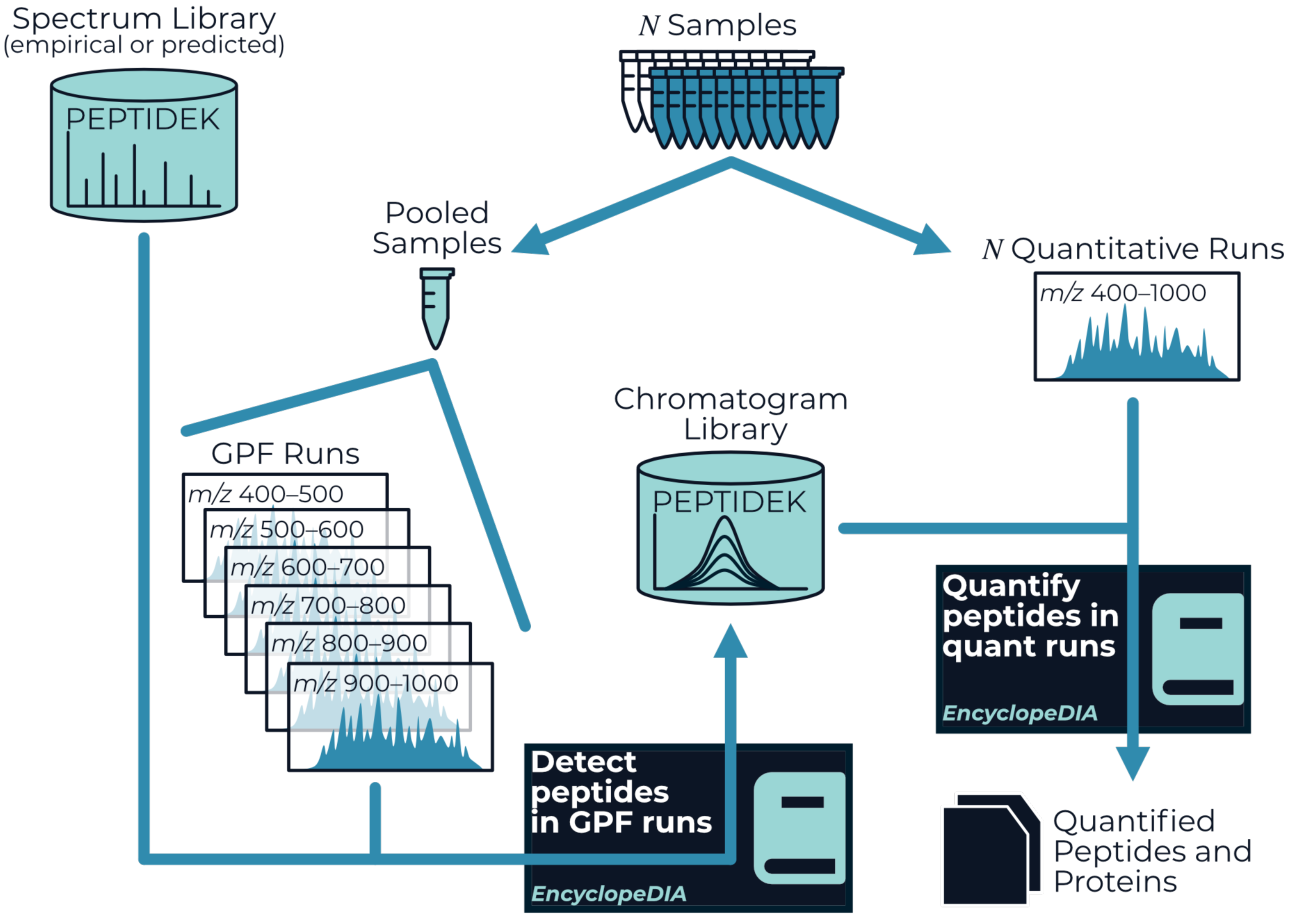


**Supplementary Figure S1: The GPF chromatogram library workflow.** Samples of interest are pooled and a chromatogram library is acquired through a series of gas phase fractionated injections. The chromatogram library is then used to detect peptides in the quantitative runs measuring individual samples, leveraging on-column retention times and instrument-tuned fragmentation patterns.

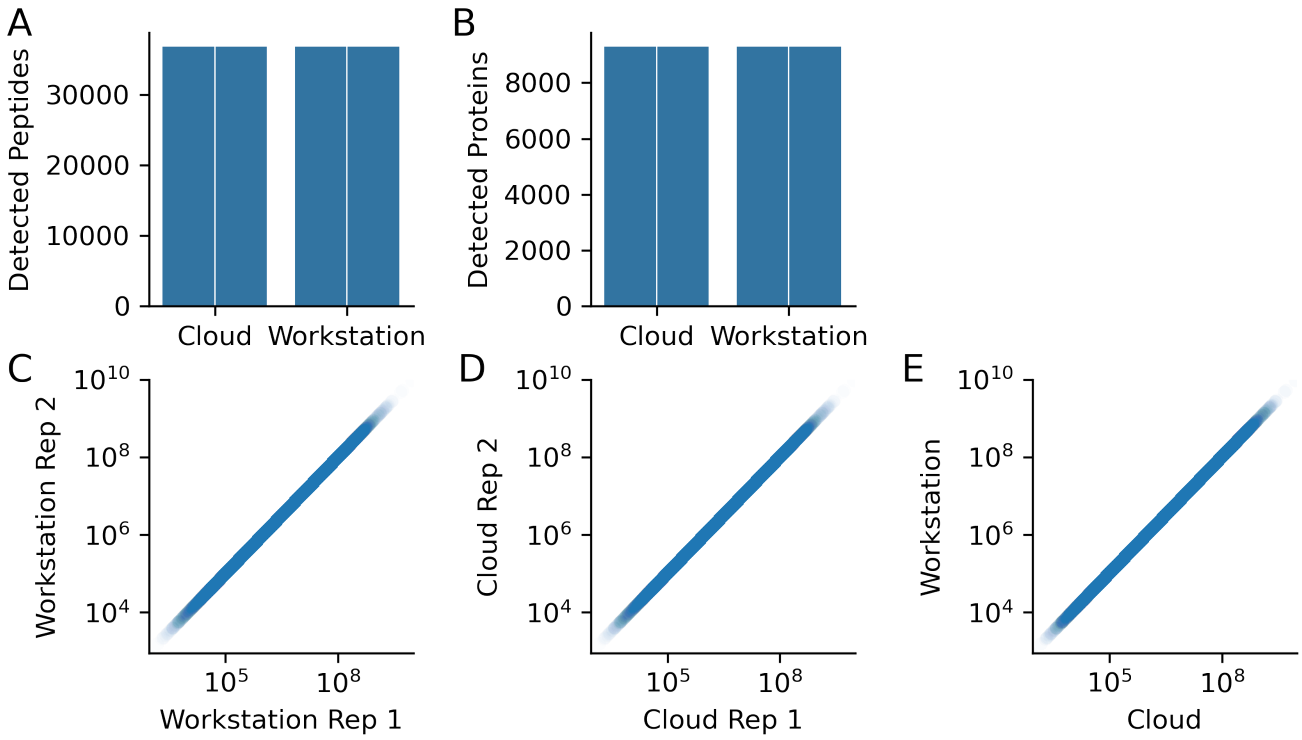


**Supplementary Figure S2: nf-encyclopedia is reproducible in the cloud and a local workstations.** Reanalysis of a three HeLa lysate injections using a six-fraction GPF chromatogram library yielded identical peptide (A) and protein (B) detections whether executed on AWS Batch or a local workstation. The peptide and protein abundances are identical between duplicate analyses (C) on a local workstation, (D) in the cloud, or (E) between the cloud and a local workstation.

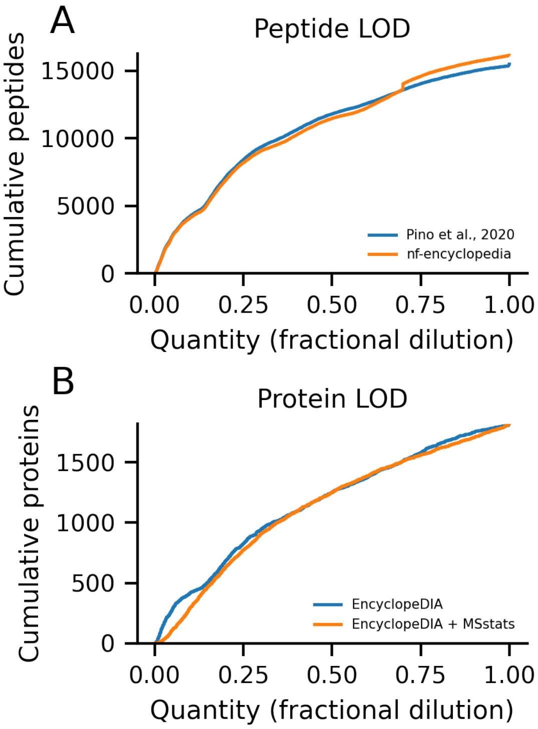


**Supplementary Figure S3: MSstats yields slightly poorer protein-level LODs than EncyclopeDIA alone.** (A) The peptide-level LODs from nf-encyclopedia closely align with the originally published analysis. (B) Protein summarization with MSstats results in higher LODs when compared with EncyclopeDIA alone.
